# Wheat pollen uptake of CRISPR/Cas9 RNP-PDMAEMA nanoassemblies results in targeted loss of gene function in progeny

**DOI:** 10.1101/2022.06.02.494465

**Authors:** Neelam Gogoi, Mehwish Kanwal, Michael Norman, Jacob Downs, Nabil Ahmad, Rohit Mago, Harbans Bariana, Markus Müllner, Urmil Bansal, Brian Jones

## Abstract

The utility of CRISPR in plants has remained limited by the dual difficulties of delivering the molecular machinery to target cells and the use of somatic cell techniques that require tissue culture-based *de novo* organogenesis. We developed 5-10 nm isodiametric polyplex nanoassemblies, comprising poly [2-(dimethylamino)ethylmethacrylate] PDMAEMA (PD) polycationic linear homopolymers and CRISPR/Cas9 ribonucleoproteins (RNPs), that enable endocytosis-driven RNP uptake into pollen grains. Pollen from wheat plants (genotype Gladius+Sr50), homozygous for monogenic *Sr50*-mediated resistance to stem rust (*Puccinia graminis* f. sp. tritici -Pgt), were incubated with RNP/PD nanoassemblies targeting the dominant, *Sr50* rust resistance gene. The treated pollen grains were then used to fertilize Gladius+Sr50 florets and the resulting M1 plants were tested for loss of *Sr50* function via rust resistance screens. The identification of fully susceptible M1 seedlings indicated that the *Sr50* RNPs acted on both alleles, indicating they were transferred via the treated pollen to the zygote. The ability to readily deliver CRISPR RNPs to reproductive cells via biodegradable, polymeric nanocomplexes has significant implications for the efficiency of gene editing in plants.

## INTRODUCTION

Plant cells are surrounded by a complex polysaccharide cell-wall matrix that acts as a barrier to exogenous materials, with a size exclusion limit (SEL) of ∼5-20 nm (1). The transfection of DNA, proteins and other materials into plant cells generally requires the use of cell wall penetrating techniques, such as biolistics (2) or *Agrobacterium tumifaciens*-mediated transformation (3), or alternatively, the creation of protoplasts via the enzymic removal of the cell wall (4). Although *in vitro* synthesized CRISPR/Cas9 ribonucleoproteins (RNPs) have been successfully transfected into plant cells (5,6), genetic modification using the above techniques generally relies on the use of somatic cells and the subsequent regeneration of whole plants via tissue culture-based *de novo* organogenesis. Most commercially important crop plants are recalcitrant to these methods (7).

Introducing CRISPR edits via germline cells offers the possibility of circumventing the use of somatic cell techniques. One of the distinctive advantages of the model dicotyledonous plant, *Arabidopsis thaliana*, is that the egg cells can be readily genetically modified *in planta* via the Agrobacterium-mediated floral dip method (8). In other species, the relative inaccessibility of the egg cells prohibits such methods. By contrast, the sperm cells are contained within the generally readily accessible pollen grains. In flowering plants, the sperm cells are delivered to the ovary via a pollen tube emanating from the germinated pollen grain. The fusion of sperm and egg cells in the ovary results in a zygote and ultimately the embryo and seed. The growth of pollen tubes towards the ovary is supported by the uptake of essential materials from the surrounding female transmitting tract via an active endocytosis system (9). S-RNAse ribonucleases, for example, are taken up into pollen tubes to participate in the regulation of pollen tube growth (10). We hypothesized that packaging RNPs into nanocomplexes would facilitate their endocytosis-driven uptake into hydrated pollen grains and that once inside they would have the capacity to access the sperm cells and potentially to be able to be transferred to the zygote during karyogamy.

Common systems for the delivery of CRISPR/Cas9 into target cells include electroporation (11), microinjection (12), viral-based delivery (13,14) and the use of nanomaterials. A multitude of nanomaterials-based approaches have been developed for mammalian systems (15-18). Polymeric nanoparticles (NPs) in particular have the advantages of a high cargo loading capacity, flexibility in terms of molecular architecture, the capacity for proton-sponge driven escape from endosomes, and are generally low-cost, and can be designed to have low levels of cytotoxicity (19,20). In plants, the cell wall has limited the use of nanomaterials-based techniques, but microparticle bombardment (21) and magnetic field attraction (22), and materials such as mesoporous silica NPs (MSNs) (23), calcium phosphate NPs (24), dendrimers (25) and surface-functionalized single-walled carbon nanotubes (SWCNTs) (26,27) have recently been shown to have some capacity to overcome the cell wall barrier. Nanomaterials-based approaches for introducing CRISPR/Cas9 to plant cells, however, have not yet been developed.

CRISPR/Cas9 can be introduced into cells as plasmid DNA, mRNA/sgRNA combinations, or directly as RNPs (28). The use of RNPs avoids issues associated with inefficient *in vivo* transcription and translation, and the potential for the unintended integration of plasmid DNA into the target cell genome (29). We investigated the possibility that a polycationic linear homopolymer, poly [2-(dimethylamino)ethylmethacrylate], PDMAEMA (PD) could facilitate RNP uptake into pollen grains as a route for introducing DNA edits directly into reproductive cells *in vivo*. To test this hypothesis, we targeted the Sr50-mediated resistance to stem rust (*Puccinia graminis* f. sp. *tritici* -Pgt) fungal infection in the rust resistant wheat (*Triticum aestivum*) genotype, Gladius+Sr50. The resistance to rust infection in Gladius+Sr50 is dependent on the monogenic presence of the dominant stem rust resistance gene, *Sr50* (*RGA1-A*). *Sr50* and several highly homologous sequences (*RGA1-B* to *RGA1-G*) (Supplementary Information) were introduced into wheat via the introgression of a fragment of the rye (*Secale cereale*) chromosome (1R) (30). ATTO550 fluorescence-labelled crRNA:tracrRNA duplex gRNAs (hereafter referred to as gRNAs) were designed to simultaneously target sequences in *Sr50* and its homologs in order to cause loss-of-function deletions, leading to the loss of rust resistance.

When combined, the RNPs and PD polymers formed ∼5-10 nm diameter spherical polyplexes that were able to be taken up into hydrated wheat pollen grains. Once internalized, the RNPs were shown to localize to the sperm cells. M1 progeny of RNP/PD-treated pollen from Gladius+Sr50 plants homozygous for *Sr50* crossed with homozygous *Sr50* Gladius+Sr50 female parents demonstrated a loss of *Sr50*-dependent rust resistance. This loss of rust resistance in the M1 plants indicates a loss of both the male and female alleles and therefore that the RNPs survived intact through fertilization to function in the zygote. The loss of rust resistance was similarly observed in the M2 progeny of self-fertilized M1 plants. Delivering RNP nanocomplexes via pollen uptake has significant implications for the integration of *de novo* gene edits in plants. The ability to apply the technique to a major crop species indicates its potential value for trait tailoring in plant breeding programs.

## RESULTS AND DISCUSSION

### Fabrication of nanoscale RNP/PD nanocomplexes

The length and structure of a polymeric chain affects its overall charge and the final size and amount of RNP that can be incorporated in the assembled nanocomplex. To test the effect of PD chain length on RNP complex formation, we synthesized two different lengths of linear PD with different molecular weights (MW) using atom transfer radical polymerisation (ATRP): PDMAEMA_82_ (PD_82_, MW∼12.8 kDa) and PDMAEMA_530_ (PD_530_, MW ∼83.2 kDa) (Supplementary Information). The MW of a polymer is directly related to its transfection efficiency and cytotoxicity. PDs >112 kDa are known to induce high levels of cytotoxicity (31). RNPs have a heterogeneous surface with an overall net negative charge (20,32) that enables them to readily associate with polycationic molecules like PD. PDs with considerable variation in MW (PD_82_ and PD_530_) were synthesized to determine whether they formed suitably sized and uniform nanoassemblies with the RNPs. The PDs were initially mixed with the RNPs at a 10 nM concentration in a 1:1 molar ratio in wheat pollen germination media (WPGM) (see Methods). As *ex vivo* hydrated wheat pollen grains can quickly lose viability, the WPGM was specifically designed to trigger and support metabolic activity in the hydrated pollen grains, but to suppress the germination process during the RNP/PD incubation period (33). Electrostatic interactions between the two components (Fig. 1a) led to the formation of RNP/PD nanoassemblies for both PD_82_ and PD_530_ (Fig. 1b & 1c). A particle distribution plot (Fig. 1d) showed that PD_82_ consistently yielded smaller assembled particulates with 90% of the particles in the ∼5-10 nm size, while PD_530_ gave more polydisperse assemblies outside this range. The surface charge analysis of nanoassemblies through zeta-potential measurements revealed a change in the charge of the RNPs from an initial - 10 mV to -0.42 mV in the RNP/PD_82_ and +1.23 mV in the RNP/PD_530_, indicating successful electrostatic interactions between the weakly negatively charged RNPs and the positively charged PDs. The 1:1 molar ratio of RNP to PD_530_ appeared to elicit coacervate agglomeration of the nanoassemblies (34,35) (Fig. 1c, inset), whereas RNP/PD_82_ resulted in a more stable assembly of relatively uniform monodisperse nanocomplexes.

**Fig. 1.**
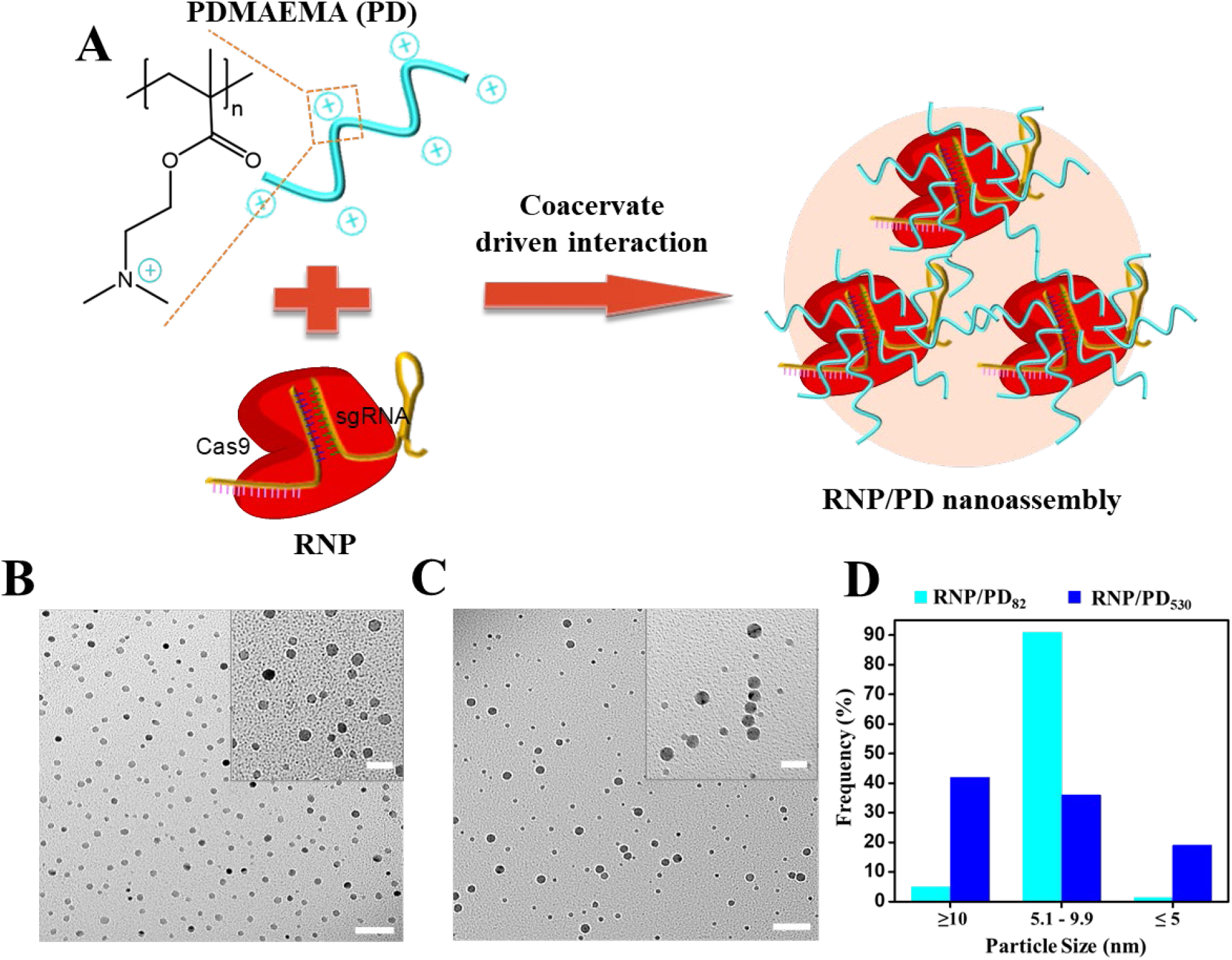
Fabrication and characterization of RNP nanocomplexes with PD of different molecular weights. **(A)** Representation of assembly of RNP/PD nanocomplexes. **(B)** TEM image of RNP/PD_82_. **(C)** TEM image of RNP/PD_530_. Scale bar, 50 nm. Inset scale bar, 20nm. **(D)** Particle size distribution quantification of TEM images.

### Delivery of RNPs into wheat pollen grains

To pre-test the capacity for the uptake of the nanocomplexes into hydrated pollen grains, RNPs were assembled with the Cas9 endonuclease and fluorescently (ATTO550) (IDT) labelled crRNA:tracrRNA duplex guide RNAs (gRNAs) (Methods and Supplementary Information). RNP/PD_82_ and RNP/PD_530_ nanocomplexes were assembled and incubated in WPGM with freshly harvested pollen from stage Z6.5-Z7.6 Gladius+Sr50 wheat spikelets (36). To determine the optimal duration of the incubation period, aliquots of freshly prepared RNP/PD_82_ and RNP/PD_530_ nanoassembly solutions were incubated for 1h, 4h, 8h in the dark at 22 °C. The efficiencies of transfection into pollen grains were determined by examining ATTO550 labelling (λ_ex_=560 nm) using a Leica SP5 laser scanning confocal microscope, with the percentage of ATTO550 fluorescence in the pollen grains determined via ImageJ analysis of the confocal images (Methods). Pollen grains treated with RNP/PD_82_ (Fig. 2a & 2b) at 1h incubation showed an overall 20% increase in ATTO550 fluorescence intensity over the control zero ATTO550 treatment (Fig. 2c). I ATTO550 fluorescence intensity increased to 41% and 74% at 4h and 8h, respectively (Fig. 2a). A similar increasing trend in ATTO550 fluorescence was observed over time in the RNP/PD_530_ treatment (Fig. 2a & 2b), with 33% at 1h, 52% at 4h and 81% at 8h. By 4h of incubation, the ATTO550-RNP fluorescence signal localized at focal points within the pollen grains (Fig. 3a). Overlaying the ATTO500-RNP fluorescence signal with DAPI DNA staining showed a co-localization of the signals, indicating that the ATTO550-labelled RNPs were associated with the sperm cell nuclei and pollen cell vegetative nucleus (Fig. 3b). At 8h of incubation with RNP/PD_530_ there was an obvious agglomeration of fluorescent particles in the solution, similar to the agglomeration observed in TEM images of the RNP/PD_530_ nanoassemblies (Fig. 1c). In the absence of the nanocomplexes in a control treatment, an average of ∼60% of pollen grains germinated in vitro by 8 hours of incubation in WPGM. In the presence of the RNP/PD_530_ nanocomplexes this was reduced to 29%. By contrast, 56% of the pollen grains treated with the RNP/PD_82_ nanocomplexes germinated by 8 hours (Supplementary Information). As a result of this and that it has been observed previously that the longer the polymer, the greater its potential for toxicity (31), the shorter chain length RNP/PD_82_ nanocomplexes were used for all subsequent experiments.

**Fig. 2.**
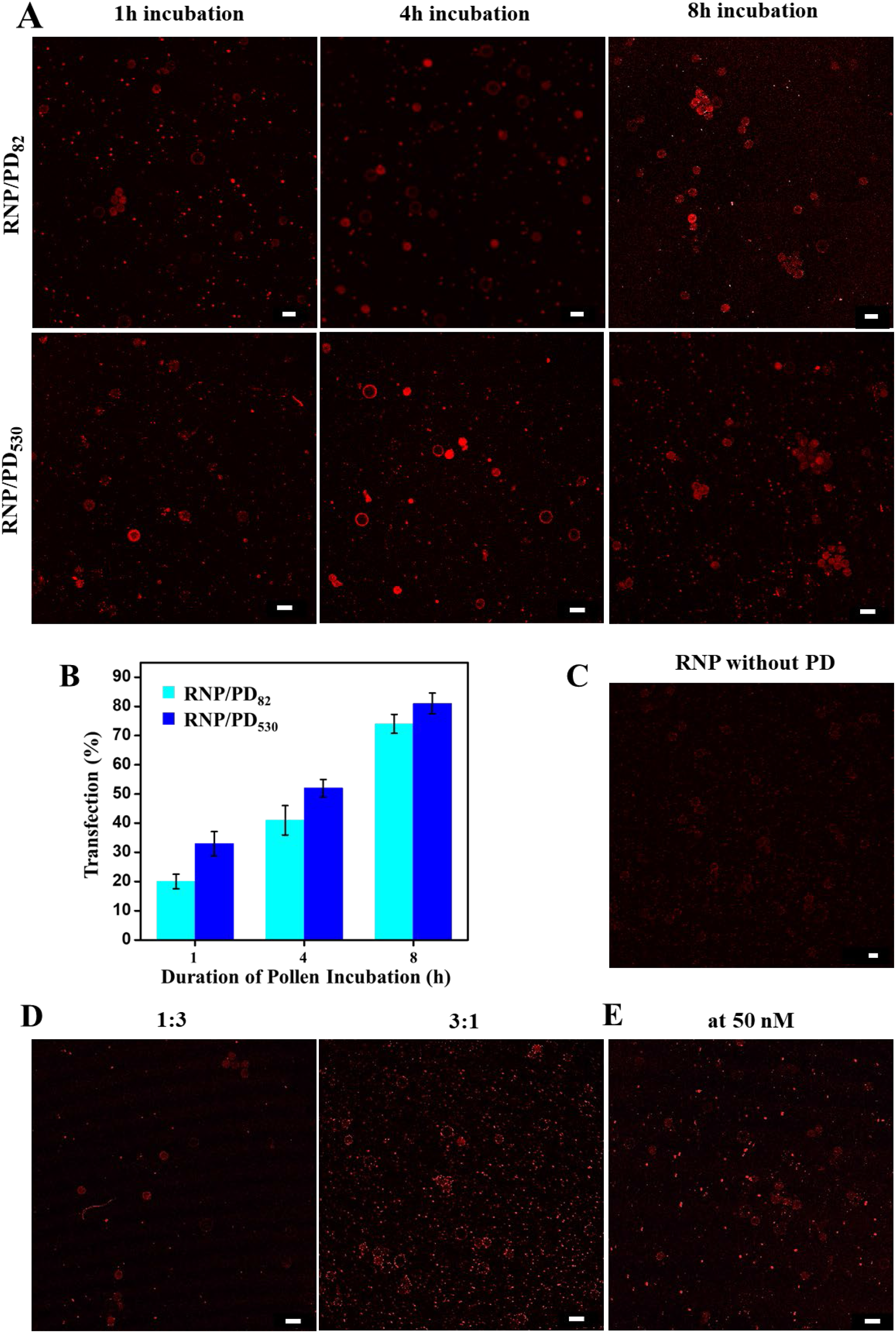
Delivery of ATTO550 labelled RNP into sperm cells in wheat pollen. **(A)** Confocal microscopy images of wheat pollen treated with RNP/PD_82_ and RNP/PD_530_ nanocomplexes followed by incubation for 1h, 4h, 8h. **(B)** Percentage of cytoplasmic/nuclear transfection efficiency of ATTO550-labelled RNPs quantified from confocal microscopy images. **(C)** Unassembled RNP observed at 8h post treatment of incubated wheat pollen grains. **(D)** Delivery of RNP/PD_82_ nanocomplexes fabricated with different molar ratios of RNPs and PD_82_, viz., 1:3 and 3:1, respectively. **(E)** Delivery of RNP/PD_82_ nanocomplexes at 50 nM of RNP and PD_82_. Scale bars, 50 μm. λ_excitation_, 560 nm. Confocal images are representative images from 5 technical replicates per 5 biological replicates per sample.

**Fig 3.**
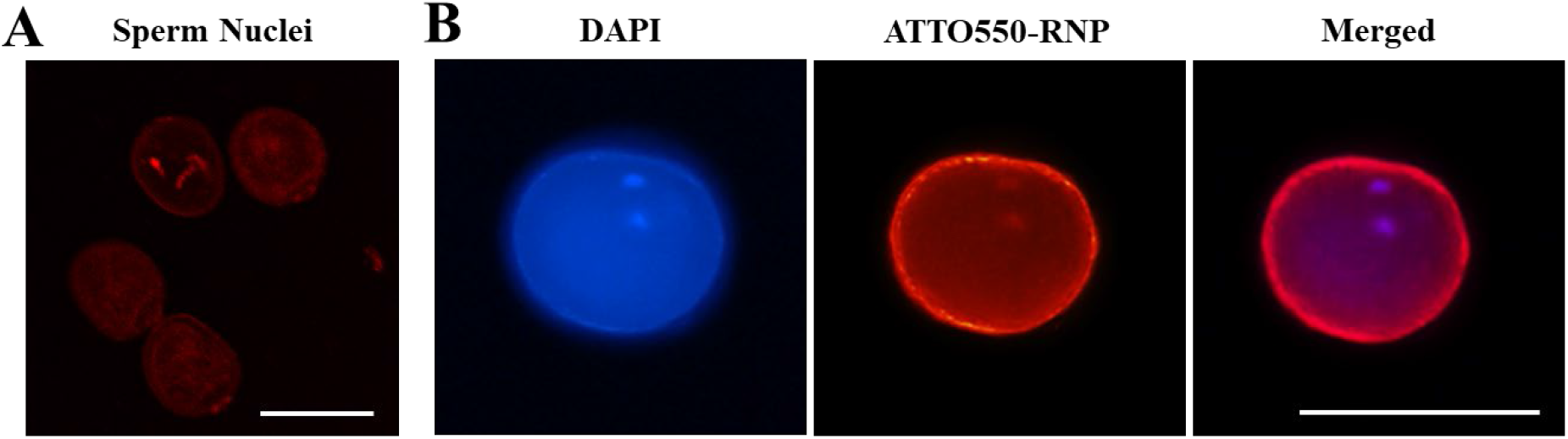
Delivery of ATTO550 labelled RNPs into sperm nuclei inside wheat pollen grains. **(A)** Confocal microscopy images showing fluorescence of concentrated ATTO550-labelled RNPs in sperm nuclei/vegetative nucleus in wheat pollen grains. **(B)** Overlapping DAPI-staining and ATTO550 RNP signals in sperm nuclei/vegetative nucleus in a wheat pollen grain. Scale bars, 10 μm.

The ratio of RNP to polymer used in the assembly of nanocomplexes has previously been shown to affect gene editing efficiencies (37). In addition to the 1:1 molar ratio, we tested RNP/PD_82_ nanocomplexes fabricated using molar ratios of 1:3 or 3:1 RNP to PD for their capacity to be incorporated into pollen grains (see Methods). At 4h of incubation with a 1:3 molar ratio of RNP/PD_82_ (Fig. 2d), there was an approximately 30% reduction in fluorescence intensity compared to the 1:1 molar ratio formulation. An agglomeration of the nanocomplex was observed on the surface of pollen grains incubated with 3:1 molar ratio of RNP/PD_82_ (Fig. 2d) or a high concentration (50 nM) of RNPs (Fig. 2e). It is possible that agglomeration reduces nanoassembly uptake. As a result, we concluded that a nanoassembly concentration of 10 nM and a 1:1 molar ratio formulation was the most suitable for both transfection efficiency and post-incubation pollen viability.

### Mechanism of RNP delivery by nanocomplexes

Nanoparticles and their cargo have been shown to internalize in mammalian cells via endocytosis (38). Wheat pollen grains have an aperture in the external wall (exine) of ∼5 μm diameter (Fig. 4a) through which the pollen tube emerges during germination. Following germination, the pollen tube grows via tip growth, requiring a constant flow of materials and active endo-and exocytosis. Both clathrin-mediated (CM) (39) and clathrin-independent (40) endocytosis pathways contribute to pollen tube tip growth. To examine the role of endocytosis in RNP/PD_82_ nanocomplexes uptake, we pre-treated treated hydrated pre-germinated pollen grains for 1h with each of chemicals that have previously been shown to act as inhibitors of endocytosis in plant cells; wortmannin (41) and ES9-17, a chemical analogue of endosidin9 (ES9) (42), prior to incubation with the RNP/PD_82_ nanocomplexes. A marked decrease in the ATTO550 fluorescence intensity was observed in pollen grains pre-treated with the inhibitors compared to the no inhibitor controls (Fig. 4b). Pre-treatment with ES9-17 led to a 73% decrease in fluorescence and the Wortmannin pre-treatment reduced fluorescence by 58% (Fig. 4c). These levels of inhibition are similar to those previously observed with the use of these inhibitors in plant cells (41,42). The stronger inhibition induced by ES9-17 suggests that clathrin-mediated (CM) endocytosis is the primary pathway for the uptake of the RNP/PD_82_ nanocomplexes into pre-germinated pollen. We hypothesize that the pathway for RNP/PD nanocomplex uptake into pollen occurred via the free movement of the complexes to the inner pollen cell wall (intine) at the exine pore site (Fig. 4a), where because they are smaller than the SEL they were able to access the pollen cell plasma membrane. A thermodynamically favourable interaction between the nanocomplexes and the plasma membrane surface is expected to have led to membrane wrapping, the formation of vesicles and the transfer of the nanocomplexes into the pollen grain cytoplasm (38,43). Once inside the cytoplasm, cationic polymers like PD are known to execute endosomal escape through a proton-sponge mechanism in order to release the cargo into the cytoplasm (31). The mechanism underlying the transfer of RNPs into sperm cells and zygote and whether the RNPs were released from the nanocomplexes prior to, or after transfer remains unclear. A full exploration of these elements of the process may lead to improvements in the technique and assist with adapting the process for other plant species.

**Fig. 4.**
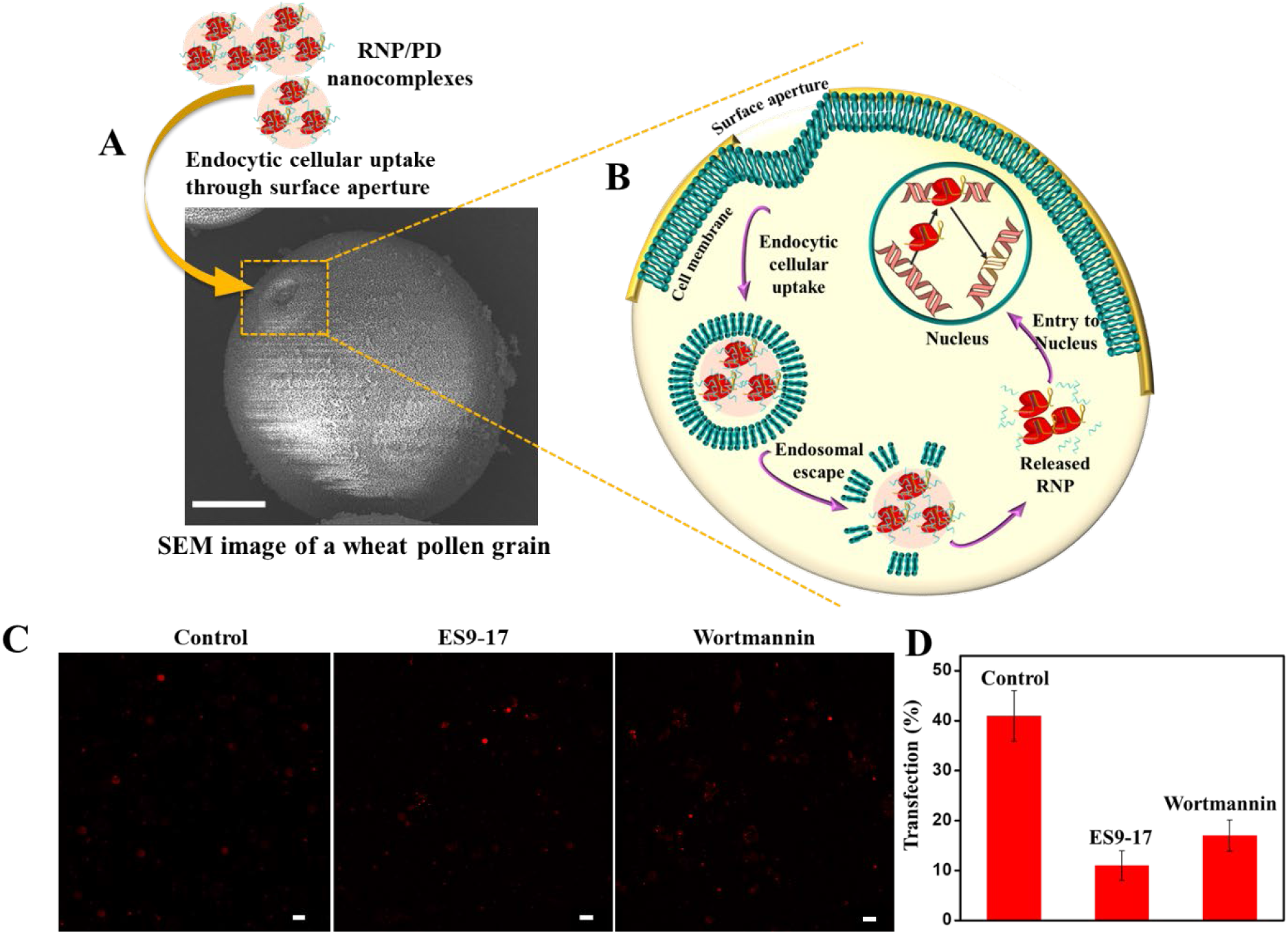
Endocytic delivery of RNP/PD nanocomplex in wheat pollen grains. **(A)** SEM image of wheat pollen showing exine surface aperture. Scale bar, 10 μm. **(B)** A schematic representation of the endocytic pathway followed by RNP/PD nanocomplexes through the aperture in the pollen grain exine. **(C)** Confocal microscopy images of wheat pollen grains pre-treated with endocytosis inhibitors; ES9-17 and Wortmannin followed by treatment with RNP/PD_82_ nanoassemblies and incubation for 4h. Scale bars, 50 μm. **(D)** Comparison of transfection (%) of ATTO550-RNP into wheat pollen grains in the presence of endocytosis inhibitors and control (ATTO-550 RNP/PD in the absence of endocytosis inhibitors).

### CRISPR/Cas9-induced mutations in wheat plants

The stem rust resistance gene, *Sr50*, confers monogenic resistance to the obligate biotrophic fungal pathogen, Pgt, that causes stem rust in wheat. The germination of stem rust spores on the leaves of susceptible wheat cultivars (e.g. Morocco) leads to the colonisation of the leaf by fungal hyphae and the production of rust-colored uredia on the leaf surface (Fig. 5a). By contrast, in Gladius+Sr50 and other genotypes carrying *Sr50*, plant-pathogen recognition events involving *Sr50* and the Pgt avirulence protein, AvrSr50, lead to localized leaf cell apoptosis that arrests the rust lifecycle prior to fungal sporulation (44). A loss of *Sr50* function leads to the loss of resistance (45).

**Fig. 5.**
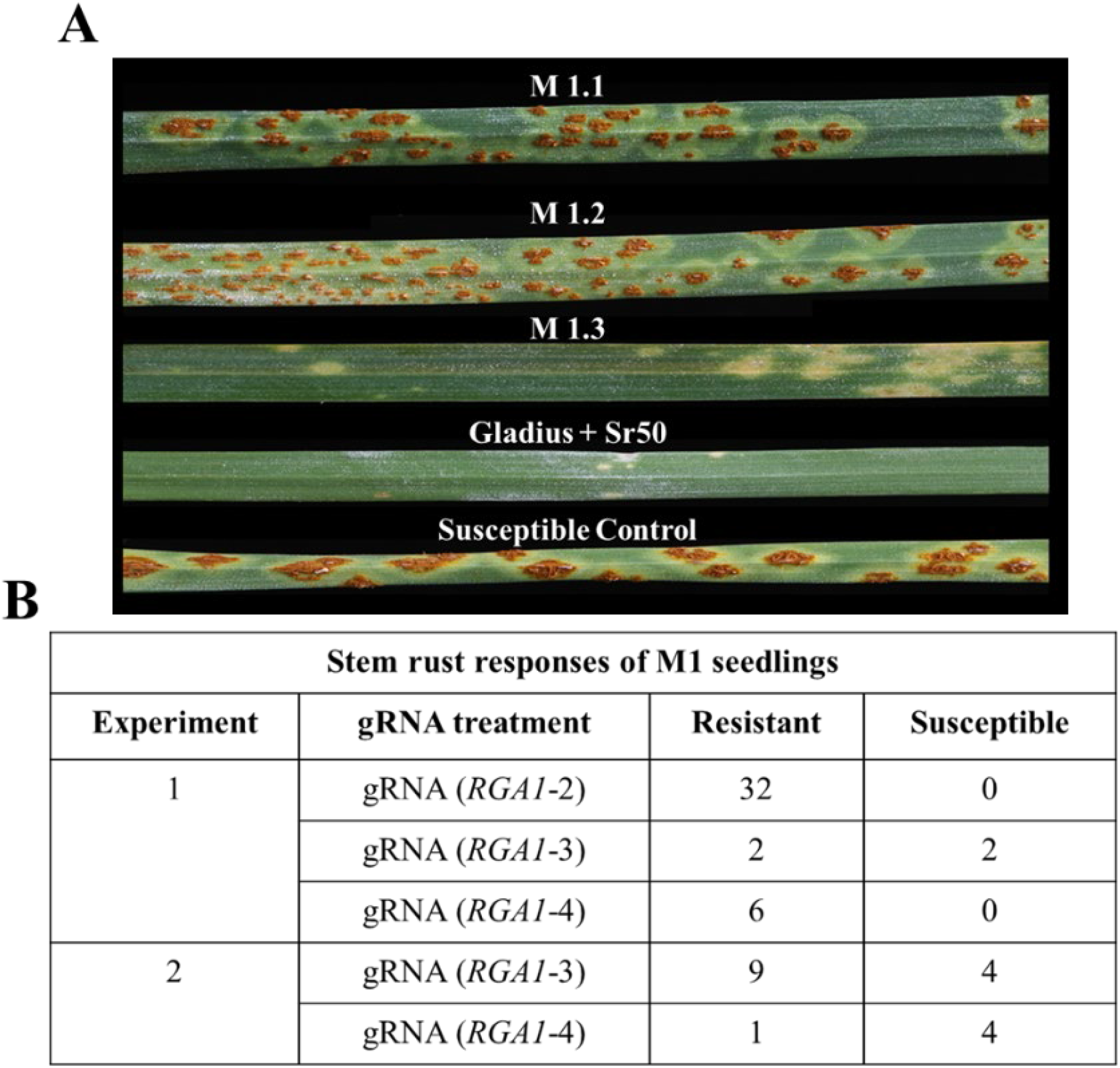
Stem rust infection phenotypes in wheat leaves. (**A)** Rust pustule development 14 days after inoculation of Gladius+Sr50 seedlings with Pgt pathotype 34-1,2,3,4,5,6,7. S1.26, S1.27 and R1.28 are leaves of M1 progeny of gRNA-2 RNP-treated Gladius+Sr50 pollen x Gladius+Sr50. The *Sr50* parent is Gladius+Sr50, and the susceptible parent is the wheat cultivar, Morocco. **(B)** Results of stem rust responses of M1 seedlings against *Sr50* avirulent Pgt pathotype 34-1,2,3,4,5,6,7.

To induce rust susceptibility in Gladius+Sr50 plants, three gRNAs; gRNA-1, gRNA-2, and gRNA-3, were designed to target *Sr50* and RGA1-B to RGA1-G homologs on the introgressed rye chromosomal fragment (30) (Supplementary Information). Sites corresponding to the gRNA sequences are also present in various endogenous wheat sequences (Supplementary Information). Simultaneously targeting *Sr50* and adjacent homologs was expected to lead to the deletion or inversion of intervening chromosomal sequences, increasing the potential for the disruption of *Sr50*-dependent rust resistance in affected progeny. The efficacy of the gRNAs was initially tested *in vitro*, using RNPs and *Sr50* PCR amplicons. All three gRNAs were shown to cut the *Sr50* cDNAs *in vitro* (Supplementary Information), indicating a potential for functionality *in planta*.

As *Sr50* has dominant inheritance, a loss of resistance to Pgt requires the loss of both alleles. In the first round of *in planta* experiments, a total of 35 M1 seeds were set in Gladius+Sr50 spikelets fertilized with RNP/PD-treated Gladius+Sr50 pollen (Fig. 5b). To assess the potential for biallelic loss of *Sr50* function, all 35 M1 seedlings were screened for susceptibility to the *Sr50*-avirulent Pgt pathotype 34-1,2,3,4,5,6,7. Thirty-three of the M1 plants retained Pgt resistance (Fig. 5a). Two of the M1 plants (S1.26 and S1.27), however, had fully susceptible phenotypes (Supplementary Information), with infection type 3+ (Fig. 5a). The loss of resistance in the M1 generation indicated bi-allelic loss of function and, hence the ability of the RNPs to survive intact through pollen tube growth, sperm cell release and karyogamy.

In experiment two, a total of 16 viable seeds were obtained from pollen grains treated with gRNA-2 and gRNA-3. Rust screening of all 16 M1 plants revealed that 11 of the seedlings retained full rust resistance. The remaining 5 seedlings were either completely susceptible to the *Sr50*-avirulent Pgt pathotype 34-1,2,3,4,5,6,7 or had levels of uredial development not seen on genotypes carrying *Sr50* (Supplementary Information). The infection types ranged from 3 to 3+ (Fig. 5a).

All 51 M1 plants from the round one and two experiments were self-fertilized to obtain M2 progeny in order to identify the presence of heterozygous loss-of-function in the M1 population. Eight to 10 seeds were planted from each M2 population and all plants were screened for stem rust response with the *Sr50*-avirulent pathotype Pgt 34-1,2,3,4,5,6,7. All progeny showed phenotypes similar to their respective M1 parent (Supplementary Information), revealing a lack of heterozygosity in the M1 plants and strongly suggesting that the CRISPR/Cas9 was preferentially active in the zygotes. Multiple studies have previously demonstrated that CRISPR/Cas9 is active in plant zygotes (46).

### Genotyping loss-of-function mutants

Oligonucleotide primers were designed to amplify sequences encompassing the gRNA-2 and gRNA-3 target sites in the *Sr50* gene to identify induced casual mutations in the susceptible lines. Genomic DNA was extracted from leaf tissue obtained from untreated control plants and from susceptible and resistant M1 plants from experiment one. The primer set 2-3 F/R (Supplementary Information) specifically targeted a sequence in *Sr50* encompassing both the gRNA-2 and the gRNA-3 target sites. The 2-3 F/R primers amplified this target *Sr50* sequence from genomic DNA extracted from untreated Gladius+Sr50 plants and M1 plants that had retained resistance to rust infection. The same 2-3 F/R primer set, however, invariably failed to amplify the target *Sr50* sequences from the M1 plants that had lost resistance to the fungus. Additional primer sets were designed to interrogate sequences upstream and downstream from the *Sr50* gRNA target sites (Supplementary Information). These primer sets similarly amplified sequences corresponding to *Sr50* from untreated plants and Pgt resistant M1 plants, but once again failed to amplify the *Sr50* sequences in the susceptible M1 plants. Although CRISPR/Cas9 can induce small indel mutations at target sites in plant zygotes (47), CRISPR-induced double-stranded DNA breaks (DSBs) can lead to large sequence deletions and chromosomal rearrangements (6). We used gRNAs targeting multiple sequences on the same introgressed rye DNA fragment in order to cause the loss of *Sr50* and therefore Sr50-mediated rust resistance. It remains to be determined if large deletions/chromosomal rearrangements are intrinsic to this method or whether unique gRNA target sites will allow for greater editing precision.

The impact of CRISPR/Cas9 and associated toolkit on plant breeding continues to expand. But the reliance on the decades-old techniques of explant somatic cell transformation and tissue culture regeneration have remained significant roadblocks. We have shown here that complexing RNPs with cationic PDMAEMA polymers enables their incorporation into viable wheat pollen grains via endocytosis. The results suggest that the RNPs remain inactive until they are incorporated in the zygote. To the best of our knowledge this is the first demonstration of the uptake of protein-polymer nanocomplexes into pollen grains. Both of the linear homopolymers trialed formed populations of spherical nanocomplexes with RNPs under the SEL predicted to allow their passage through plant cell walls. The uniformity, colloidal stability and negligible cytotoxicity of the smaller RNP/PD82 nanocomplexes indicates their suitability for *in planta* delivery. Further experimentation may identify polymer architectures and compositions with improved *in planta* efficiencies. Although the mechanisms involved in the delivery of the RNPs to the sperm cells and zygote remain to be elucidated, the work successfully demonstrates that this facile nanoparticle delivery mechanism holds considerable potential for accelerating CRISPR/Cas9 gene editing and crop improvement.

## MATERIALS AND METHODS

### Materials

All chemical reagents were used as received without further purification unless specified otherwise. 2-(dimethylamino) ethyl methacrylate (DMAEMA, ≥98%), 1,1,4,7,10,10-hexamethyltriethylenetetramine (HMTETA, ≥97%), ethyl α-bromoisobutyrate (EB*i*B, ≥98%), 1,4-dioxane (≥99%), 2-(N-Morpholino)ethanesulfonic acid hydrate (MES hydrate, C_6_H_13_NO_4_S.xH_2_O, ≥99.5%), magnesium chloride (MgCl_2_, ≥98%), boric acid (H_3_BO_3_, ≥99.9%), calcium nitrate tetrahydrate (Ca(NO_3_)_2_.4H2O, ≥99%), magnesium sulphate (MgSO_4_, ≥98%), sucrose (≥99.7 %), polyethylene glycol 4000 (PEG4000) were purchased from Sigma-Aldrich. Copper(I) chloride (CuCl, ≥97%), acetone (≥99%) hexane (≥99%), anisole (≥99%) were ordered from Merck. DMAEMA was passed through a column of aluminium oxide to remove the inhibitor. Integrated DNA Technologies (IDT, Skokie, IL) provided recombinant *Streptococcus pyrogenes* CRISPR Cas9 nuclease V3 (62 μM, MW= 162400 g/mol) containing nuclear localisation sequences (NLS) and a C-terminal 6-His tag obtained from the purification of an *E. coli* strain expressing the nuclease. The crRNAs (10 nmol, MW= 11,715g/mol) and ATTO550-labelled tracrRNA (20 nmol, MW= 22,937.4 g/mol) were also provided by IDT.

### Methods

#### Preparation of Wheat Pollen Germination Media (WPGM)

Wheat Pollen Germination media (WPGM) was used to prepare and characterise the nanoassemblies in all experiments. The WPGM was prepared by mixing boric acid (0.01 g), CaNO_3_ (0.2 g), MgSO_4_ (0.2 g), sucrose (0.8 M, 5 g) in 100 mL Milli Q water under continuous stirring using a magnetic stir bar in a Schott bottle until a clear solution is obtained. PEG4000 (10 g) was then slowly added to the solution and allowed to mix well for 5-10 min. Finally, pH of the pollen germination media was adjusted to 6.5 (34).

#### Assembly of RNPs

For the assembly of RNPs, the crRNAs were first hybridized with equimolar concentrations of ATTO550-labelled tracrRNA (IDT) to obtain ATTO550-labelled crRNA:tracrRNA duplexes (gRNA). The gRNAs were then assembled with Cas9 (IDT) at a 1:1 molar ratio. See supplementary information for further details.

#### Nanocomplexing of RNPs with PD

A stock solution of 1 μM concentration of PD_82_ and PD_530_ was prepared in MES buffer at pH 6. RNP was aliquoted out to make a stock of 1 μM concentration in Milli Q water. The 1 μM stock solutions of PD_82_ and PD_530_ were dispersed in freshly prepared WPGM to achieve a 10 nM concentration. An equimolar concentration of preassembled RNP was mixed with the PD-WPGM solution to prepare a 1:1 molar ratio formulation of nanocomplexes. For example, to prepare a 200 μL solution of nanocomplex, 2 μL of 1 μM PD solution was dispersed in 196 μL of WPGM and vortexed for 30 sec. This PD-WPGM solution was added drop-by-drop to 2 μL of 1 μM preassembled RNP in a separate vial under a continuous stirring condition for 45 min. The mixture was incubated for at least 30 min at 22 °C for the formation of the nanocomplexes. A fresh solution of nanocomplexes should be prepared just prior to pollen treatment in order to avoid the aggregation of particles. A 1:1 molar ratio nanocomplex at 50 nM was prepared following the same protocol as described above, by adjusting the net concentration of PD and RNP to 50 nM.

For the 1:3 RNP-PD_82_, 10 nM of RNP was assembled with 30 nM of PD-WPGM solution under constant stirring. Similarly, 3:1 RNP-PD_82_ was prepared by slowly mixing 30 nM RNPs with 10 nM of the PD-WPGM solution under constant stirring. Both solutions were incubated at 22 °C for 30 min before use.

#### Plant growth and collection of wheat pollen grains

The homozygous wheat genotype Gladius+Sr50 was produced through backcrossing in the Australian Cereal Rust Control Program (H.S. Bariana unpublished results). See Supplementary Information for details.

#### Confirmation of delivery of RNP nanocomplex into wheat pollen grains with laser scanning confocal microscopy fluorescence intensity analysis

For pollen treatments, a 100 μL aliquot of the RNP/PD nanocomplex solution was added to a glass microscope slide. Pollen grains were added into the nanocomplex solution with forceps and mixed delicately to achieve a uniform spread. The glass slide was then covered with a 24×60 mm coverslip and sealed to incubate the samples in the dark for 1h to 8h (for a time course experiment; 1h, 4h, 8h). Five biological replicates were prepared for each nanocomplex treatment. A Leica SP5 II Confocal and Multiphoton microscope was used to detect the presence of the ATTO550 labelled RNPs in the wheat pollen. Samples were analysed with 560 nm excitation and 50% laser power at 10X magnification with pinhole = 70.6 μm for the emission range of 566 nm to 630 nm at a resolution of 12 bit. At least 5 non-overlapping confocal fields of view of each replicate were saved. ImageJ analysis was conducted in order to obtain the mean value of fluorescence for every pollen grain in a confocal field of view for each replicate. These measurements were averaged to obtain a mean value of fluorescence intensity correlating to the internalization percentage of each treatment.

### Statistical analysis

#### Confocal data: ATTO550 labelled RNP delivery into wheat pollen grains

N=5 technical replicates, i.e, 5 non-overlapping confocal fields of view per sample were imaged for 5 independent replicates of each treatment. The % transfection value was calculated through the grayscale value of each pollen grain using ImageJ. A mean value was obtained for every independent image of a treatment. The final percentage transfection is expressed as a mean value ± SD obtained from mean values of each independent image per treatment.

### TEM data

Particle size distribution of the RNP/PD_82_ and RNP/PD_530_ nanocomplexes was assessed through grayscale analysis in ImageJ for 150 particles. For every 30 particles per treatment, a Student’s *t* test was applied and any difference with *P* <0.05 was considered significant.

### *In planta* treatment

For the *in planta* experiments, freshly harvested Gladius+Sr50 pollen grains were incubated for 1 hour in WPGM with 10 nM 1:1 RNP/PD_82_ nanocomplexes, containing either gRNA-1, gRNA-2 or gRNA*-*3 (Supplementary Information). Immediately post treatment, the hydrated pollen grains were transferred to the stigmatic surface of emasculated Z6.5-Z7.6 Gladius+Sr50 flowers for pollination and fertilization. Crossing bags were used to cover the pollinated flowers to avoid unintentional cross-fertilization events.

### DNA extraction, PCR conditions and sequencing

Gladius+Sr50 plants were grown in 9-cm diameter pots. Twenty grams of complete fertilizer Aquasol was dissolved in 9L of water was applied before and after sowing. DNA was extracted from each sample following the methodology in Bansal and co-workers (45). Two more applications of Aquasol were applied when seedlings were 14 and 21 days old. For the rust resistance experiments, seedlings were inoculated at the two-leaf stage, with the Pgt pathotype 34-1,2,3,4,5,6,7 by brushing a mixture of urediniospores and talcum powder on the leaf adaxial surface. Inoculated seedlings were then incubated in a dew chamber for 48 h before being moved into a greenhouse room maintained at 25 °C. Rust response assessments were made 14-16 days post-inoculation using the scale described in McIntosh and co-workers (48).

Each PCR reaction contained 60 ng of wheat genomic DNA (30 ng/μL), 1 μL of Immolase DNA polymerase buffer, 1 μL of DNTPs, 0.25 μL of each primer, 0.04 μL of Immolase DNA Polymerase and 5.5 μL of milli Q water. The PCRs were performed on a CFX100 under the following cycling conditions: 10 mins at 95 °C; five touchdown cycles of 30 s at 94 °C, 30 s at 65-60 °C (dropped by 1 °C per cycle), 30 s at 72 °C; and 30-35 cycles of 30 s at 94 °C, 30 s at 65-60 °C, 30 s at 72 °C; with a final 5 mins extension at 72°C for 5 mins. PCR products were run on a 2% agarose gel in 1% TBE (Tris-borate-EDTA) buffer with 2-3 μL of gel red/100 mL of gel solution. PCR products to be sequenced were eluted using the Bioline Isolate II PCR and Gel Kit to manufacturer’s instructions. Sanger sequencing was performed by the Australian Genome Research Facility (AGRF).

## Acknowledgments

We thank Wenjun Gu, Kiran Soni, Wenkai Lyu, Fahad Ali, Bede Johnston, Hanif Miah, Lincoln Hoang-linh Ngo, the members of the University of Sydney, Polymer Nanochemistry Lab, KCPC, School of Chemistry group and the University of Sydney Microscope and Microanalysis (SMM) facility for their contributions.

## Author contributions

Conceptualization: BJ, MM, UB, HB

Methodology: NG, MM, UB, BJ

Investigation: NG, MK, MN, JD, UB, RN

Visualization: NG

Supervision: BJ, MM, UB, HB

Writing: NG, BJ, MM, UB, HB

N.G† and M.K† contributed equally to the work described in the manuscript.

## REFERENCES AND NOTES

1. G. S. Demirer, H. Zhang, N. S. Goh, R. L. Pinals, R. Chang, M. P. Landry. Carbon nanocarriers deliver siRNA to intact plant cells for efficient gene knockdown. Sci. Adv. 6, eaaz0495 (2020).

2. A. Hayashimoto, Z. Li, N. Murai, A Polyethylene Glycol-Mediated Protoplast Transformation System for Production of Fertile Transgenic Rice Plants. Plant Physiol. 93, 857–863 (1990).

3. S.N. Char, A.K. Neelakandan, H. Nahampun, B. Frame, M. Main, M. H. Spalding, P. W. Becraft, B. C. Meyers, V. Walbot, K. Wang, B. Yang, An Agrobacterium-delivered CRISPR/Cas9 system for high-frequency targeted mutagenesis in maize. Plant biotechnol. J. 15, 257–268 (2017).

4. H. Hamada, Y. Liu, Y. Nagira, R. Miki, N. Taoka, R. Imai, Biolistic-delivery-based transient CRISPR/Cas9 expression enables in planta genome editing in wheat. Sci. Rep. 8, 1–7 (2018).

5. J.W. Woo, J. Kim, S.I. Kwon, C. Corvalán, S.W. Cho, H. Kim, S.G. Kim, S.T. Kim, S. Choe, J.S. Kim, DNA-free genome editing in plants with preassembled CRISPR-Cas9 ribonucleoproteins. Nat. Biotechnol., 33, 1162–1164 (2015).

6. Z. Liang, K. Chen, T. Li, Y. Zhang, Y. Wang, Q. Zhao, J. Liu, H. Zhang, C. Liu, Y. Ran, C. Gao, Efficient DNA-free genome editing of bread wheat using CRISPR/Cas9 ribonucleoprotein complexes. Nat. Commun. 8, 1–5 (2017).

7. L. Herrera-Estrella, J. Simpson, M. Martínez-Trujillo, “[Transgenic Plants, An Historical Perspective]” in Transgenic Plants: Methods and Protocols ((Springer Science & Business Media., ed. 286, 2005), pp. 3–31.

8. S. J. Clough, A.F. Bent, Floral dip: a simplified method for Agrobacterium-mediated transformation of Arabidopsis thaliana. Plant J. 16, 735–743 (1998).

9. G. Grebnev, M. Ntefidou, B. Kost, Secretion and endocytosis in pollen tubes: models of tip growth in the spot light. Front. Plant Sci. 8, 154, (2017).

10. D.T. Luu, X. Qin, D. Morse, M. Cappadocia, S-RNase uptake by compatible pollen tubes in gametophytic self-incompatibility. Nature 407, 649–651 (2000).

11. M. Zuckermann, V. Hovestadt, C.B. Knobbe-Thomsen, M. Zapatka, P.A. Northcott, K. Schramm, J. Belic, D.T. Jones, B. Tschida, B. Moriarity, D. Largaespada, Somatic CRISPR/Cas9-mediated tumour suppressor disruption enables versatile brain tumour modelling. Nat Commun. 6, 1–9 (2015).

12. T. Sakuma, S. Nakade, Y. Sakane, K.I.T. Suzuki, T. Yamamoto, MMEJ-assisted gene knock-in using TALENs and CRISPR-Cas9 with the PITCh systems. Nat. Protoc. 11, 118–133 (2016).

13. D. Wang, P.W. Tai, J. Riley, G. Gao, J.A. Rivera-Pérez, Streamlined ex vivo and in vivo genome editing in mouse embryos using recombinant adeno-associated viruses. Nat. Commun. 9, 1–12 (2018).

14. R.J. Platt, S. Chen, Y. Zhou, M.J. Yim, L. Swiech, H.R. Kempton, J.E. Dahlman, O. Parnas, T.M. Eisenhaure, M. Jovanovic, D.B. Graham, CRISPR-Cas9 knockin mice for genome editing and cancer modelling. Cell 159, 440–455 (2014).

15. J.A. Zuris, D.B. Thompson, Y. Shu, J.P. Guilinger, J.L. Bessen, J.H. Hu, M.L. Maeder, J.K. Joung, Z.Y. Chen, D.R. Liu, Cationic lipid-mediated delivery of proteins enables efficient protein-based genome editing in vitro and in vivo. Nat. Biotechnol. 33, 73–80 (2015).

16. B.T. Staahl, M. Benekareddy, C. Coulon-Bainier, A.A. Banfal, S.N. Floor, J.K. Sabo, C. Urnes, G.A. Munares, A. Ghosh, J.A. Doudna, Efficient genome editing in the mouse brain by local delivery of engineered Cas9 ribonucleoprotein complexes. Nat. Biotechnol. 35, 431–434 (2017).

17. R. Mout, M. Ray, G. Yesilbag Tonga, Y.W. Lee, T. Tay, K. Sasaki, V.M. Rotello, Direct cytosolic delivery of CRISPR/Cas9-ribonucleoprotein for efficient gene editing. ACS Nano 11, 2452–2458 (2014).

18. H. Zhu, L. Zhang, S. Tong, C.M. Lee, H. Deshmukh, G. Bao, Spatial control of in vivo CRISPR-Cas9 genome editing via nanomagnets. Nat. Biomed. Eng., 3, 126–136 (2019).

19. Y. Gong, S. Tian, Y. Xuan, S. Zhang, Lipid and polymer mediated CRISPR/Cas9 gene editing. J. Mater. Chem. B 8, 4369–4386 (2020).

20. G. Chen, A.A. Abdeen, Y. Wang, P.K. Shahi, S. Robertson, R. Xie, M. Suzuki, B.R. Pattnaik, K. Saha, S. Gong, A biodegradable nanocapsule delivers a Cas9 ribonucleoprotein complex for in vivo genome editing. Nat Nanotechnol. 14, 974–980 (2019).

21. S. Martin-Ortigosa, J.S. Valenstein, V.S.-Y. Lin, B.G. Trewyn, K. Wang, Gold Functionalized Mesoporous Silica Nanoparticle Mediated Protein and DNA Codelivery to Plant Cells Via the Biolistic Method. Adv. Funct. Mater. 22, 3576–3582 (2012).

22. Y. Hao, X. Yang, Y. Shi, S. Song, J. Xing, J. Marowitch, J. Chen, J. Chen, Magnetic gold nanoparticles as a vehicle for fluorescein isothiocyanate and DNA delivery into plant cells. Botany 91, 457–466 (2013).

23. F.P. Chang, L.Y. Kuang, C.A. Huang, W.N. Jane, Y. Hung, C.H. Yue-ie, C.Y. Mou, A simple plant gene delivery system using mesoporous silica nanoparticles as carriers. J Mater. Chem. B 1, 5279–5287 (2013).

24. S. Naqvi, A.N. Maitra, M.Z. Abdin, M.D. Akmal, I. Arora, and M.D. Samim, Calcium phosphate nanoparticle mediated genetic transformation in plants. J Mater. Chem. 22, 500–3507 (2012).

25. L. Jiang, L. Ding, B. He, J. Shen, Z. Xu, M. Yin, X. Zhang, Systemic gene silencing in plants triggered by fluorescent nanoparticle-delivered double-stranded RNA. Nanoscale 6, 9965–9969 (2014).

26. G.S. Demirer, H. Zhang, J.L. Matos, N.S. Goh, F.J. Cunningham, Y. Sung, R. Chang, A.J. Aditham, L. Chio, M.J. Cho, B. Staskawicz, High aspect ratio nanomaterials enable delivery of functional genetic material without DNA integration in mature plants. Nat. Nanotechnol. 14, 456–464 (2019).

27. S.Y. Kwak, T.T.S. Lew, C.J. Sweeney, V.B. Koman, M.H. Wong, K. Bohmert-Tatarev, K.D. Snell, J.S. Seo, N.H. Chua, M.S. Strano, Chloroplast-selective gene delivery and expression in planta using chitosan-complexed single-walled carbon nanotube carriers. Nature nanotechnol. 14, 447–455 (2019).

28. H.X. Wang, M. Li, C.M. Lee, S. Chakraborty, H.W. Kim, G. Bao, K.W. Leong, CRISPR/Cas9-based genome editing for disease modeling and therapy: challenges and opportunities for nonviral delivery. Chem. rev. 117, 9874–9906 (2017).

29. X. Ma, Q. Zhu, Y. Chen, Y.G. Liu, CRISPR/Cas9 platforms for genome editing in plants: developments and applications. Mol. plant 9, 961–974 (2016).

30. R. Mago, P. Zhang, S. Vautrin, H., Šimková, U. Bansal, M.C. Luo, M. Rouse, H. Karaoglu, S. Periyannan, J. Kolmer, Y. Jin, The wheat Sr50 gene reveals rich diversity at a cereal disease resistance locus. Nat. Plants 1, 1–3 (2015).

31. S. Agarwal, Y. Zhang, S. Maji, A. Greiner, PDMAEMA based gene delivery materials. Mater. Today 15, 388–393 (2012).

32. Z. Glass, M. Lee, Y. Li, Q. Xu, Engineering the delivery system for CRISPR-based genome editing. Trends Biotechnol. 36, 173–185 (2018).

33. M. Kanwal, Optimisation of in vitro techniques for pollen-mediated gene editing in wheat (Triticum aestivum L). Plant Reprod. (2020) (In-press).

34. C.E. Sing, S.L. Perry, Recent progress in the science of complex coacervation. Soft Matter 16, 2885–2914 (2020).

35. S.N. Warnakulasuriya, M.T. Nickerson, Review on plant protein-polysaccharide complex coacervation, and the functionality and applicability of formed complexes. J. Sci. Food Agric. 98, 5559–5571 (2018).

36. J.C. Zadocks, T.T. Chang, C.F. Konzak, The growth stage code for cereals. Weed Res. 14, 415–421 (1974).

37. Z. Tan, Y. Jiang, M.S. Ganewatta, R. Kumar, A. Keith, K. Twaroski, T. Pengo, J. Tolar, T.P. Lodge, T.M. Reineke, Block Polymer Micelles Enable CRISPR/Cas9 Ribonucleoprotein Delivery: Physicochemical Properties Affect Packaging Mechanisms and Gene Editing Efficiency. Macromol. 52, 8197–8206 (2019).

38. S. Behzadi, V. Serpooshan, W. Tao, M.A. Hamaly, M.Y. Alkawareek, E.C. Dreaden, D. Brown, A.M Alkilany, O.C. Farokhzad, M. Mahmoudi, Cellular uptake of nanoparticles: journey inside the cell. Chem. Soc. Rev. 46, 4218–4244 (2017).

39. J. Derksen, T. Rutten, I.K., Lichtscheidl, A.H.N. De Win, E.S. Pierson, G. Rongen, Quantitative analysis of the distribution of organelles in tobacco pollen tubes: implications for exocytosis and endocytosis, Protoplasma 188, 267–276 (1995).

40. A. Moscatelli, Distinct endocytic pathways identified in tobacco pollen tubes using charged nanogold J. Cell Sci. 120, 3804–3819 (2007).

41. Q. Liu, B. Chen, Q. Wang, X. Shi, Z. Xiao, J. Lin, X. Fang, Carbon nanotubes as molecular transporters for walled plant cells. Nano Lett. 9, 1007–1010 (2014).

42. W. Dejonghe, I. Sharma, B. Denoo, S. De Munck, Q. Lu, K. Mishev, H. Bulut, E. Mylle, R. De Rycke, M. Vasileva, D.V. Savatin, Disruption of endocytosis through chemical inhibition of clathrin heavy chain function. Nat. Chem. Bio. 15, 641–649 (2019).

43. P. Foroozandeh, A.A. Aziz, Insight into cellular uptake and intracellular trafficking of nanoparticles. Nanoscale Res. Lett., 13, 1–12 (2018).

44. J. Chen, N.M. Upadhyaya, D. Ortiz, J. Sperschneider, F. Li, C. Bouton, S. Breen, C. Dong, B. Xu, X. Zhang, R. Mago, Loss of AvrSr50 by somatic exchange in stem rust leads to virulence for Sr50 resistance in wheat. Science 358, 1607–1610 (2017).

45. U. K. Bansal, A. G. Kazi, B. Singh, R. A. Hare, H. S. Bariana, Mapping of durable stripe rust resistance in a durum wheat cultivar Wollaroi. Mol. Breed 33, 51–59 (2014).

46. A. Korablev, V. Lukyanchikova, I. Serova, N. Battulin, On-Target CRISPR/Cas9 Activity Can Cause Undesigned Large Deletion in Mouse Zygotes. Int. J. Mol. Sci.. 21, 3604 (2020).

47. E. Toda, T. Okamoto, CRISPR/Cas9-Based Genome Editing Using Rice Zygotes. Curr Protoc Plant Biol. 5, e20111 (2020).

48. R.A. McIntosh, C.R. Wellings, R.F. Park, in Wheat Rusts: An Atlas of Resistance Genes. (CSIRO Publishing, 1995).

49. Cordeiro, R. A. et al. Synthesis of well-defined poly(2-(dimethylamino)ethyl methacrylate) under mild conditions and its co-polymers with cholesterol and PEG using Fe(0)/Cu(II) based SARA ATRP, Polym. Chem. 4, 3088–3097 (2013).

50. Zhu, T. et al. Optical maps refine the bread wheat Triticum aestivum cv. Chinese Spring genome assembly. Plant J. 107, 1, 303–314 (2021).

51. Corpet, F. Multiple sequence alignment with hierarchical clustering, 1988, Nucl. Acids Res., 6, 10881–10890.

52. Bariana, H. & McIntosh, R. Cytogentic studies in wheat. XV. Location of rust resistance genes in VPM1 and their genetic linkage with other disease resistance genes in chromosome 2A. Genome / National Research Council Canada = Génome / Conseil national de recherches Canada, 36, 476–82 (1993).

