## Supplemental Materials for "Wheat pollen uptake of CRISPR/Cas9 RNP-PDMAEMA nanoassemblies results in targeted loss of gene function in progeny"

**This PDF file includes:**

Synthesis of polymers
Analytical techniques
Assembly of RNP
Plant growth and collection of pollen grains
Guide RNA sequences and target sites
Endocytosis study: Pre-treatment of wheat pollen with endocytosis inhibitors
Figs. S1 to S6
Tables. S1 to S2

**Synthesis of polymers**

PDMAEMA (PD) was prepared by atom transfer radical polymerisation (ATRP) (49), as shown in Fig. S1. All of the reagents and the monomer were combined in a one-neck Schlenk flask equipped with a stir bar and subjected to 3-5 freeze-pump-thaw cycles to remove oxygen from the reaction mix. The Schlenk flask was then placed into an oil bath at 65 °C with continuous stirring under nitrogen flow for 1.5 or 3 h. This was followed by rapid cooling with liquid nitrogen and exposure to air. The purified PD was obtained by 3 cycles of precipitation-centrifugation-precipitation into an excess of hexane, followed by redispersion in acetone. To obtain polymers in their dry form, acetone was evaporated using a Rotavap (Buchi), followed by freeze-drying in 1,4-dioxane. Further details on the synthesis and characterisation for each PD are set out below:

**$PDMAEMA_{82}$ ($PD_{82}$):** EBIB (3 mg, 15.38 µmol), HMTETA (3.54 mg, 15.38 µmol), CuCl (1.52 mg, 15.38 µmol), DMAEMA (241.5 mg, 153.8 µmol), anisole (0.26 mL). Reaction time=1.5h. 1H NMR conversion: 82%, Mn, NMR=12800 $gmol^{-1}$, Đ=1.23. Fig. S1.

**$PDMAEMA_{530}$ ($PD_{530}$):** EBIB (3 mg, 15.38 µmol), HMTETA (3.54 mg, 15.38 µmol), CuCl (1.52 mg, 15.38 µmol), DMAEMA (241.5 mg, 15380 µmol), anisole (2.6 mL). Reaction time=3h. 1H NMR conversion: 53%, Mn, NMR=83200 $gmol^{-1}$, Đ=1.32. Fig. S1.

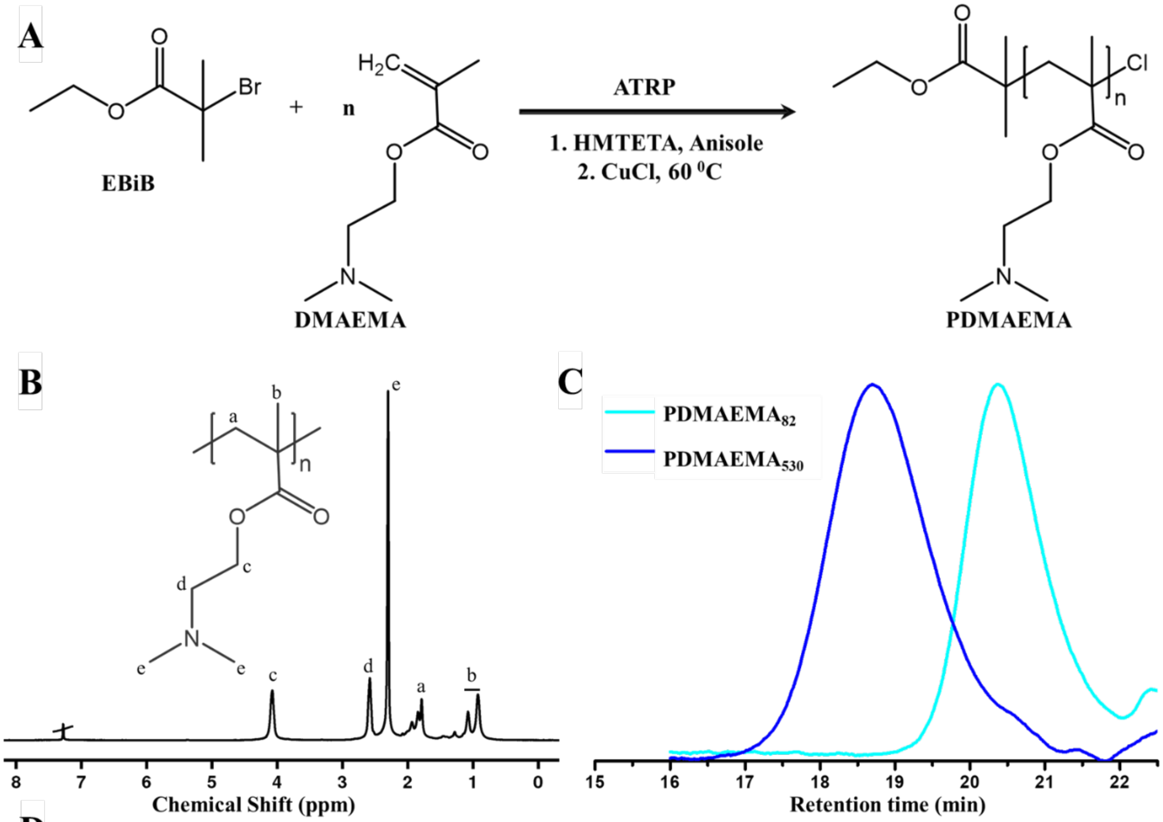


**D**

| Polymer | Reactant molar ratio [monomer] : [initiator] : catalyst | Reaction Time | DP (%) | <Mn> NMR g/mol | Dispersity (Đ) |
|---|---|---|---|---|---|
| **$PDMAEMA_{82}$** | **100:1:1** | **1.5 h** | **82** | **12800** | **1.23** |
| **$PDMAEMA_{530}$** | **1000:1:1** | **3 h** | **53** | **83200** | **1.32** |

**Fig. S1 Synthesis and characterisation of PDMAEMA (PD) of DP~ 82 and 530. (A)** Atom Transfer Radical Polymerization (ATRP) reaction of DMAEMA (D) in the presence of EB*i*B as an initiator, HMTETA as a ligand and CuCl as a catalyst in anisole. **(B)** 1H-NMR spectrum of purified PD in $CDCl_3$. **(C)** SEC (Size Exclusion Chromatograph) traces of PD of DP~ 82 and 530. **(D)** Reactant molar ratio, reaction time, degree of polymerisation (DP %), molecular weight, <Mn> NMR and dispersity (SEC) of $PD_{82}$ and $PD_{530}$.

### Analytical Techniques

The degree of polymerisation and purity of the PDMAEMA (PD) was analysed by recording proton nuclear magnetic resonance ($^1$H NMR) spectra on 300 MHz on a Bruker Advance 300 spectrometer. PD samples were prepared in deuterated chloroform ($CDCl_3$, 99%, Cambridge Isotopes Laboratories). All NMR spectra were referenced against the peak of residual reaction solvent (anisole) found in $CDCl_3$. The value of chemical shifts is reported in parts per million (ppm).

Dispersity (Đ) of the PD was determined by Size Exclusion Chromatography (SEC) measurements performed on a UFLC Shimadzu Prominence Chromatograph SEC system. *N*,*N*-dimethyl acetamide (DMAc) was used as an eluent at a flow rate of 1 mL/min and temperature of 50 °C. The polymers were dissolved in DMAc containing LiBr (0.03 % w/w) and butylated hydroxytoluene (0.05 % w/w) and transferred to a GPC vial through a 0.22 µm PTFE syringe filter before injection.

The Zeta potential measurements were performed on a Malvern Nano ZS. Freshly prepared samples were equilibrated for 2 min at 25 °C to optimise the measurements. The zeta potential values were instrumentally measured using the Smoluchowski equation.

The morphology and size of the RNP/PD nanoassemblies were visualised on a FEI Technai F12 120 kV transmission electron microscopy (TEM) in bright field mode. TEM grids (300-mesh formvar/carbon coated, ProSciTech) were glow discharged for 30 s in a Diener plasma technology system immediately before use. This process makes TEM grids hydrophilic for a better adsorption of samples. 2 µL of freshly prepared nanoassemblies was pipetted onto a TEM grid for 45 s, blotted with the edge of a filter paper, followed by washing with milli Q water three times. Similarly, a 2% solution of uranyl acetate dissolved in milli Q water was used to negatively stain the deposited sample and allowed to dry for 1 min at room temperature before they were used for the microscopic analysis. TEM images were analysed using Image J software.

**Assembly of Ribonucleoprotein (RNP) complexes**

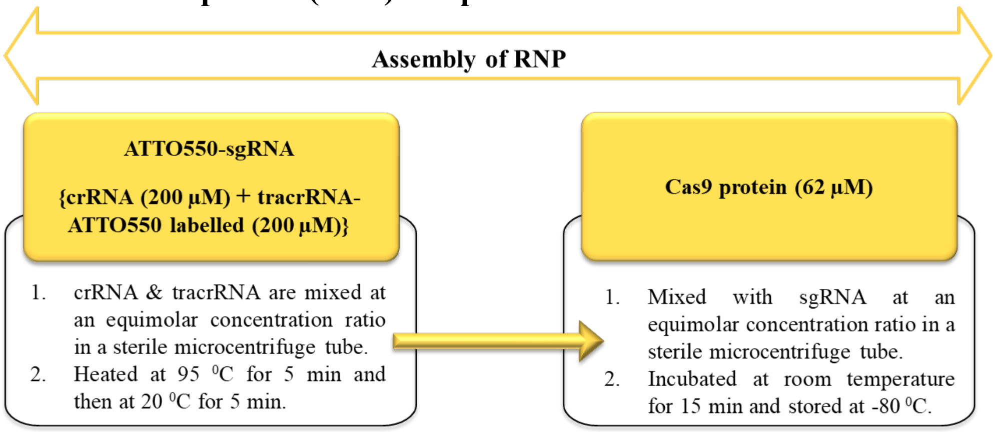


**Fig. S2 Assembly of CRISPR ribonucleoprotein (RNP) complexes.** Note: Integrated DNA Technologies (IDT) Nuclease-free buffer was used for the dilution of RNPs.

**Plant growth and collection of wheat pollen grains**

The stock used in this study was a backcross derivative of cultivar Gladius, homozygous for the introgressed fragment from *Secale cereale* chromosome 1R, carrying *Sr50* (*RGA1-A)* and homologs, *RGA1-B* to *RGA1-G* (Gladius+Sr50) (30). The plants were grown in specialised microclimate growth rooms at the University of Sydney, Plant Breeding Institute (PBI), Camden, Australia. 9-cm diameter pots filled with a soil mixture comprised of composted pine bark and sand in a 2:1 ratio mixed with fungicide-treated fertiliser (10gm per pot) were used to grow the wheat plants to flowering. Pots were then fertilised with 25g per 10 L of water Aquasol® (Hortico Pty. Ltd., Revesby, NSW, Australia) for 100 pots. Two seeds were sown in each pot to get good tiller production. Pots were then transferred to a microclimate room set at 19-20 °C and humidity 72-73% with 16 hours of day-light duration. A single dose of urea was applied to plants 10 days after sowing. Plants were raised to Z6.5-Z7.6 as per the Zadoks growth scale (36) and fresh pollen grains of Gladius+Sr50 at Z6.5-Z7.6 stage were collected from three mature anthers near to dehiscence for *in vitro* incubation with the nanoassemblies at room temperature (22 °C) in dark and humid condition. The cultivar "Morocco" was used as a susceptible control for the rust screening.

**A.**

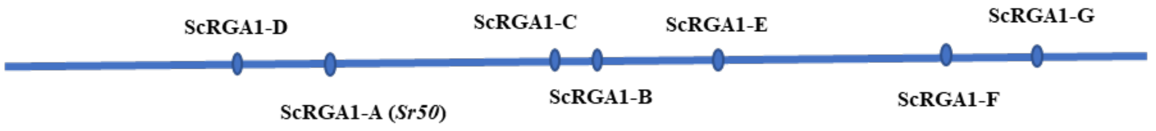


**B.**

**gRNA-1**

```
                      AATCAAGGAGCAACTCCAGGAGG
RGA1-A              GCTGACGCGATCAAGGAAATCAAGGAGCAACTCCAGGAGGTGGCTGCTAGGCGTGACAGGA
RGA1-C              GCTGACGCCATCAAGGAAATCAAGGAGCAACTCCAGGAGGTGGCTGCTAGGCGTGACAGGA
RGA1-D              GCTGACGCCGTCAAGGAAATCAAGGAGCAACTCCAGGAGGTGGCTGCTAGGCGTGACCGGA
RGA1-G              GCTGACGCTGTCAAGGAAATTAAGGAGCAACTCCAGGAGGTGGCTGCTAGGCGTGACAGGA
TraesCS1A03G0065600 GCTGACGCAGTCAAGGAAATCAAGGAGCAACTCCAGGAGGTGGCTGCTAGGCGTGACAGGA
TraesCS1B03G0068600 GCTGACGCGGTCAAGGAAATCAAGGAGCAACTCCAGGAGGTGGCTGCTAGGCGTGACAGGA
TraesCS1B03G0068700 GCTGACGCTGTCAAGGAAATCAAGGAGCAACTCCAGGAGGTGGCTGCTAGGCGTGAAAGGA
TraesCS1B03G0070700 GCTGACGCCATCAAGGAAATCAAGGAGCAACTCCAGGAGGTGTCTGCTAGGCGTGACAGGA
TraesCS1B03G0071200 GCTGACGCCATAAAGGAAATCAAGGAGCAACTCCAGGAGGTGGCTGCTAGGCGTGAAAGGA
TraesCS1B03G0072100 GCTGACGCGATCAAGGAAATCAAGGAGCAACTCCAGGAGGTGGCCGCTAGGCGTGAGAGGA
TraesCS1D03G0053300 GCTGACGCCGTCAAGGAAATCAAGGAGCAACTCCAGGAGGTGGCTGCTAGGCGTGACAGGA
```

**gRNA-2**

```
                      ATATTGTACGTCACACGGAAGGG
RGA1-A              AATAGTAGTGTGTAAATATATTGTACGTCACACGGAAGGGTTCAGAATAATTTTCAAGGTC
RGA1-G              AATAGTAGTGTGTAAATATATTGTACCTCACACGGAAGGGTTCGGAATAATTTTCAAGGTC
TraesCS1A03G0065600 AATAGTAGTGTGTAAATATATTGTACGTCACACGGAAGGGTTCAGAATAATTTTCAAGGTC
TraesCS1B03G0068600 ATCAGTTGTGTGTAAATATATTGTACGTCACACGGAAGGGTTCAGAATAATTTTCAAGGTC
```

**gRNA-3**

```
                      CCTCATTGATCTCGGCAACCCTC
RGA1-A              AAGGTTTTAAGGGATATCCTCATTGATCTCGGCAACCCTCACTCAGATCTTGCTCTTGCT
RGA1-B              AAGGTTTTAAGGGATATCCTCATTGATCTCGGCAACCCTCACTCAGACCTTGCTATACTA
RGA1-D              AAGGTTTTAAGGGATATCCTCATTGATCTCGGCAACCCTCACTCGGATCTTGCAATGCTG
RGA1-E              AAGGTTTTAAGGGATATCCTCATTGATCTCGGCAACCCTCACTCAGATCTTGCAATGCTG
RGA1-G              AAGGTTTTAAGGGATATCCTCATTGATCTCGGCAACTCTCACTCAGATCTTGCGATACTG
TraesCS1A03G0065600 AAGGTTTTAAGGGATATCCTCATTGATCTCGGCAACTCTCACTCAGATCTTGCGATACTG
TraesCS1B03G0068700 AAGGTTTTCAGGGATATCCTCATTGATCTCGGCAACTCTCACTCAGATCTGGCGATACTG
TraesCS1B03G0070700 AAGGTTTTAAGGGATATCCTCATTGATCTCGGCAACGCTCACTCAGATCTTGCGACGCTG
TraesCS1D03G0053300 AAGGTTTTAAGGGATATCCTCATTGATCTCGGCAACCCTCACTCAGATCTTGCGATACTG
```

**Fig. S3. Guide RNA (gRNA) target sites and sequences in the introgressed rye and homologous wheat sequences. A.** Introgressed genomic sequence containing the *Sr50* (*RGA1-A*) and homologs, *RGA1-B* to *RGA1-G.* **B.** gRNA-1, gRNA-2, and gRNA-3 sequences (in red text) and the associated protospacer adjacent motif (PAM) sequence (yellow highlight) identified in the *Sr50* sequence (RGA1-A). *RGA1B-RGA1G* are the corresponding sequences present in the introgressed rye chromosome 1R fragment in the Gladius+Sr50 genotype. The TraesCS1n sequences were identified by Blast analysis of the wheat cultivar, Chinese Spring (50). Corresponding sequences with more than 1 mismatch were not included in the sequence comparisons. Multiple sequence alignments were performed with Multalin (http://multalin.toulouse.inra.fr/multalin/) (51). The gRNAs were designed using the species-independent sgRNA Scorer 2.0 gRNA prediction software (https://sgrnascorer.cancer.gov/).

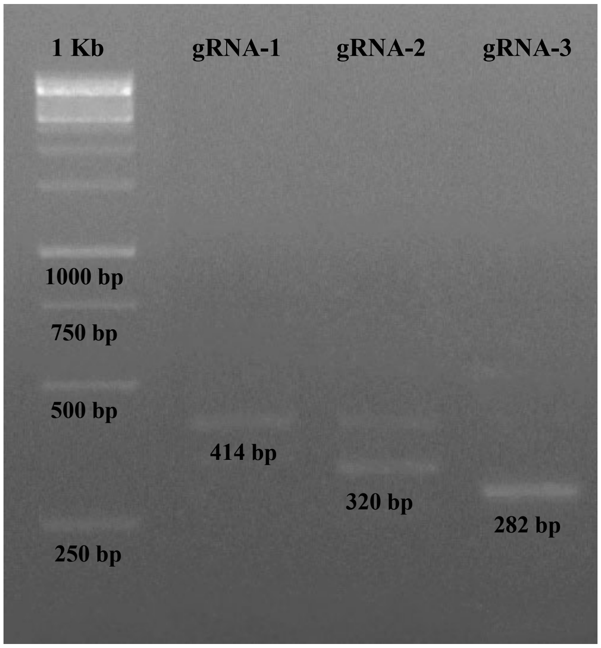


**Fig. S4. Pre-validation *in vitro* DNA cleavage test.** The RNPs used in the main experiments were initially screened for activity in vitro, targeting a PCR amplicon amplified from Gladius+Sr50 with the oligonucleotide primer set, 2-3 F/R. The RNPs incorporating gRNA-1, gRNA-2, or gRNA-3 were incubated with the PCR amplicon at 37 °C for 1 hour. Lane 1, 1kb DNA ladder. Lane 2, gRNA-1. The band shown is 414bp cleaved from a 455bp PCR amplicon. Lane 3, gRNA-2. The band shown is 320bp cleaved from a 405bp PCR amplicon. There is uncut product (405bp) indicating inefficient targeting by gRNA-2. Lane 4, gRNA-3. The band shown is 282bp cleaved from a 405bp PCR amplicon. The products were separated on a 1.5% agarose gel.

**Table S1. Responses of M1 and M2 seedlings to *Puccinia graminis* f. sp. *tritici* pathotype 34-1,2,3,4,5,6,7** (52).

| ***In planta* experiment** | **gRNA used** | **Sample** | **M1 Phenotype** | **M2 Phenotype** |
|---|---|---|---|---|
| **Round 1** | gRNA-1 | R1.1 | ; | 1- |
| | | R1.2 | ; | 1- |
| | | R1.3 | ; | 1- |
| | | R1.4 | ; | 1- |
| | | R1.5 | ; | 1= |
| | | R1.6 | ; | 1- |
| | | R1.7 | ; | 1- |
| | | R1.8 | ; | 1- |
| | | R1.9 | ; | 1- |
| | | R1.10 | ; | 0 |
| | | R1.11 | ; | 1- |
| | | R1.12 | ; | 0 |
| | | R1.13 | ; | ; |
| | | R1.14 | ; | 1- |
| | | R1.15 | ; | 1- |
| | | R1.16 | ; | 1- |
| | | R1.17 | ; | 1- |
| | | R1.18 | ; | 1- |
| | | R1.19 | ; | 1- |
| | | R1.20 | ; | 1- |
| | | R1.21 | ; | 1- |
| | | R1.22 | ; | 1- |
| | | R1.23 | ; | 1- |
| | | R1.24 | ; | 1- |
| | | R1.25 | ; | 0 |
| | gRNA-*2* | S1.26 | 3+ | 3+ |
| | | S1.27 | 3+ | 3+ |
| | | R1.28 | ; | ; |
| | | R1.29 | ; | ; |
| | gRNA-3 | R1.30 | ; | 1- |
| | | R1.31 | ; | 1- |
| | | R1.32 | ; | 1- |
| | | R1.33 | ; | 1- |
| | | R1.34 | ; | 1- |
| | | R1.35 | ; | 1- |
| **Round 2** | gRNA-2 | R2.1 | ; | ;1 |

| | | | | |
|---|---|---|---|---|
| | | R2.2 | ; | ;1 |
| | | R2.3 | ;1 | 1 |
| | | R2.4 | ;1 | 1 |
| | | R2.5 | ;1 | 1 |
| | | R2.6 | ; | ; |
| | | R2.7 | 1 | ; |
| | | R2.8 | ;1 | ; |
| | | S2.9 | 3 | 3 |
| | | R2.10 | ; | 0; |
| | | R2.11 | 1 | 1 |
| | gRNA-3 | S2.12 | 3+ | 3+ |
| | | S2.13 | 3 | 3 |
| | | S2.14 | 3+ | 3+ |
| | | S2.15 | 33+ | 33+ |
| | | R2.16 | 0; | 0; |

**Table S2. Oligonucleotide sequences used in the study.**

| Usage | Primer Name | Sequence (5'-3') |
|---|---|---|
| In vitro for gRNA 1, amplify target site | gRNA 1 F | CGATCCAGAGAGCTCATCCTCCTG |
| In vitro for gRNA 1, amplify target site | gRNA 1 R | CCTGTCACGCCTAGCAGCCA |
| In vitro for gRNA 2/3, amplify target sites | gRNA 2-3 F | GTTGGGCAAGACCACTCTTGC |
| In vitro for gRNA 2/3, amplify target sites | gRNA 2-3 R | GAAAGTGAGCACCTATAAGG |
| gRNA | gRNA 1 | AAUCAAGGAGCAACUCCAGG |
| gRNA | gRNA 2 | AUAUUGUACGUCACACGGAA |
| gRNA | gRNA 3 | GAGGGUUGCCGAGAUCAAUG |
| Amplify promoter region | For2 | AGTTGTCCAGAGCTTGGGAGACTG |
| Amplify promoter region | Rev3 | GCGAGATCTGGACCAAGCCCA |
| Amplify downstream region | 7For | TTGGCATTGGTCCAAGAAAT |
| Amplify downstream region | 7Rev | GGTGATTAGCCGACTGCCTA |

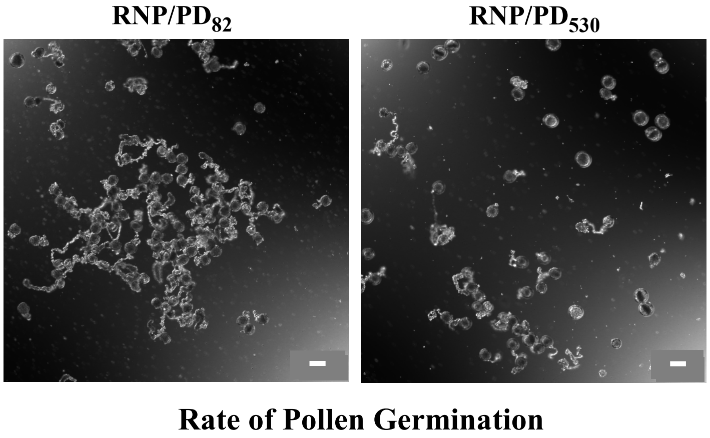


**Fig. S5. DIC Confocal Microscopy images of germination of wheat pollen.** Germination rate of wheat pollens treated with RNP/$PD_{82}$ and RNP/$PD_{530}$ at 8h post-treatment incubation. Scale bars, 50 µm.

**Endocytosis study: Pre-treatment of wheat pollen with endocytosis inhibitors**

1 mM stock solutions of the two endocytosis inhibitors; wortmannin (41) and ES9-17 (42) were prepared in DMSO. A 200 µL solution containing 30 µM of inhibitor is prepared in WPGM and 50 µL of it was added to freshly extracted wheat pollen grains for 30 min. Then 50 µL of RNP/$PD_{82}$ nanocomplex solution was added and incubated for 4 h at 20 °C in dark before examining to obtain the quantitative % fluorescence intensity under a Leica SP5 confocal microscope.

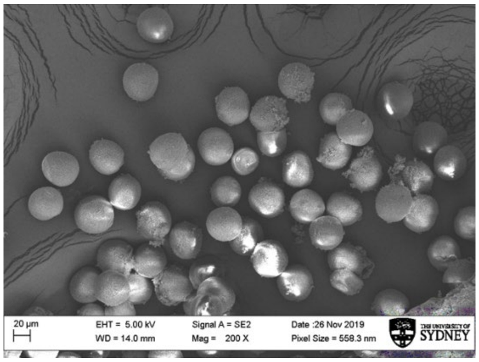


**Fig. S6. Endocytic delivery.** DIC Confocal Microscopy images of wheat pollen treated with a, RNP/$PD_{82}$ and b, RNP/$PD_{530}$ show inhibition in RNP delivery in the presence of the endocytosis inhibitors; Wortmannin and ES9-17. Scale bars, 50 µm. c, SEM image of a wheat cultivar Gladius+Sr50 pollen grains.
